# Increased H3K27 trimethylation contributes to cone survival in a mouse model of cone dystrophy

**DOI:** 10.1101/2022.02.07.479069

**Authors:** Annie L. Miller, Paula I. Fuller-Carter, Klaudija Masarini, Marijana Samardzija, Kim W. Carter, Rabab Rashwan, Alicia A. Brunet, Abha Chopra, Ramesh Ram, Christian Grimm, Marius Ueffing, Livia S. Carvalho, Dragana Trifunović

## Abstract

Inherited retinal diseases (IRDs) are a heterogeneous group of blinding disorders, which result in dysfunction or death of the light-sensing cone and rod photoreceptors. Despite individual IRDs being rare, collectively, they affect up to 1:2000 people worldwide, causing a significant socioeconomic burden, especially when cone-mediated central vision is affected. This study uses the *Pde6c*^*cpfl1*^ mouse model of achromatopsia, a cone-specific vision loss IRD, to investigate the potential gene-independent therapeutic benefits of a histone demethylase inhibitor GSK-J4 on cone cell survival. We investigated the effects of GSK-J4 treatment on cone cell survival *in vivo* and *ex vivo* and changes in cone-specific gene expression via single-cell RNA sequencing. A single intravitreal GSK-J4 injection led to transcriptional changes in pathways involved in mitochondrial dysfunction, endoplasmic reticulum stress, among other key epigenetic pathways, highlighting the complex interplay between methylation and acetylation in healthy and diseased cones. Furthermore, continuous administration of GSK-J4 in retinal explants increased cone survival. Our results suggest that IRD-affected cones respond positively to epigenetic modulation of histones, indicating the potential of this approach in the development of a broad class of novel therapies to slow cone degeneration.

## 1. Introduction

Inherited retinal diseases (IRDs) are a group of heterogeneous conditions that result in permanent death, dysfunction, or developmental delay of the cells in the retina, particularly the cone and rod photoreceptors [1]. Patients with IRD have varying levels of visual impairment, with a large proportion eventually becoming legally blind [1]. Despite their orphan designation, they collectively affect 1:2000 people worldwide [1] and pose a heavy socioeconomic burden on patients and families with an estimated cost of £523 million to the United Kingdom alone in 2019 [2]. Polls indicate most people consider blindness the worst possible ailment they could acquire [3]. Unfortunately, limited treatments are available for IRDs, with most patients relying on visual aids to enhance visual acuity, and only a small subset of people with IRD may be eligible for the only FDA-approved ocular gene therapy drug Luxturna® or visual prosthesis [4, 5]. Over the years, considerable effort has been applied in elucidating the cell death mechanisms involved in IRD pathogenesis to find novel targeting points within common cell death pathways to facilitate mutation-independent treatments. Photoreceptor degeneration in IRD was classically thought to be caused by apoptosis [6, 7] but a number of recent studies revealed a novel non-apoptotic cell death pathway [8-11]. One interesting hypothesis proposed is that high cyclic guanosine monophosphate (cGMP) content present in IRD photoreceptors is responsible for a cGMP-dependent activation of protein kinase G, which can lead to over-activation of histone deacetylases (HDAC) shown to be a major constituent of non-apoptotic cell death governing photoreceptor loss in IRD [8, 12].

HDACs are ubiquitously expressed enzymes that remove acetyl groups from histones, allowing post-translational modification, chromatin structure modulation, and changes in gene expression [13]. The opposing enzymes, histone acetyltransferases (HATs), add acetyl groups, usually leading to open chromatin and increased gene expression [13]. Histone proteins may also be modified by removing or adding other chemical groups, including methyl groups. Histone demethylases (HDM) and histone methyltransferases (HMT) work in conjunction, with HDM removing methyl groups and HMT adding methyl groups. Unlike acetylation, where adding a chemical group usually leads to an increase in transcription, histone methylation results in either gene expression or repression depending on which residue of the histone protein is methylated [14]. Some methylation examples that result in gene repression include trimethylation of H3K27 and H3K9 residues, while methylation of H3K4 and H3K36 drives active gene expression [14]. The balance between these post-translational histone modifications is essential, as numerous HDAC and HDM enzymes have functional interplay and are often concomitant, with histone demethylation also being reported as a secondary target of HDAC inhibition [15]. In the retina, HDAC inhibition suppresses cone photoreceptor cell loss in mouse models of two different types of IRDs, affecting either cones only (achromatopsia) or both rods and cones (retinitis pigmentosa, RP) [16-18]; however, there is limited knowledge on the role of histone methylation in IRD. One study has suggested that upregulation of specific histone methylation sites may be associated with disease in the *rd1* mouse model of RP, with global inhibition shown to delay rod photoreceptor degeneration and improve visual function [19]. A recent study showed that the administration of a lysine demethylase 1 inhibitor in the *rd10* model of RP prevented rod death and preserved vision [20]. However, the role of histone methylation in the degeneration of cones specifically has yet to be investigated. As cone photoreceptors are affected in most types of IRD and are responsible for our colour, acuity and daylight vision, understanding the post-translational modifications under disease conditions could be crucial in the development of novel therapies.

The present study investigates the role of H3K27me3 in cone photoreceptor cell death in a mouse model of a human type of cone dystrophy (achromatopsia). We used the *Pde6c*^*cpfl1*^ model, which shows early onset cone loss at postnatal day (PN) 14 and peak of cell death at PN24 due to mutations in the cone-specific *Pde6c* gene [21]. We highlight the relationship between methylation and acetylation by observing increased cone-specific H3K27me3 levels following treatment with the HDAC inhibitor Trichostatin A (TSA). Interestingly, the increased H3K27me3 levels in treated cones resemble the presence of H3K27me3 staining in wildtype (wt) cones. Further, we investigated the impact of GSK-J4, an H3K27 HDM inhibitor, on cone survival *in vivo* and *ex vivo*. The investigation evaluated the effect of GSK-J4 in *Pde6c*^*cpfl1*^ mutant mice via histological analysis, functional testing, and single-cell RNA sequencing. We show that a single intravitreal GSK-J4 injection generated extensive alterations in gene expression, with significant enrichment of disease-related pathways, including mitochondrial dysfunction, endoplasmic reticulum stress, and epigenetic pathways. More importantly, continuous administration of GSK-J4 to *Pde6c*^*cpfl1*^ retinal explants resulted in a 31.9% increase in cone numbers compared to sham controls, indicating the potential of epigenetic modulation for the treatment of cone loss in IRD.

## 2. Materials and Methods

### 2.1 Animals

Mice utilized in this project were housed at The University of Tübingen, University of Zürich, or The Harry Perkins Institute of Medical Research Bioresource Facility. Animals were provided with either 12/12 or 14/10 hour light/dark cycle and *ad libitum* access to food and water, with experiments performed in accordance with the ARVO Statement for the Use of Animals in Ophthalmic and Vision Research, the regulations of the Tübingen University committee on animal protection, veterinary authorities of Kanton Zürich and The Harry Perkins Institute of Medical Research’s animal ethics committee. The *Pde6c*^*cpfl1*^ mouse line (B6.CXB1-*Pde6c*^*cpfl1*^/J), originally from the Jackson Laboratory (Strain #003678) [22, 23], was used for *in vivo* Trichostatin A applications and for the *ex vivo* work. C57BL/6J mice (Jackson Laboratory, Bar Harbor Maine, USA) served as the wildtype (wt) controls. For *in vivo* GSK-J4 treatment, the *Pde6c*^*cpfl1*^ line was crossbred with the Chrnb4.EGFP line (Mutant Mouse Research Resource Centre Stock #000259, herein referred to as Chrnb4.GFP) to specifically label the cones with GFP [24, 25]. Consequently, this line is referred to as *Pde6c*.GFP. Chrnb4.GFP mice on a C57BL/6J background were used as controls for *in vivo* GSK-J4 work.

### 2.2 Ex vivo retinal explant culture

Organotypic retinal cultures (including retinal pigment epithelium, RPE) were prepared under sterile conditions from *cpfl1* (*n =* 13) and wildtype (*n =* 4) animals, as previously described [16, 17]. PN14 animals were euthanized by cervical dislocation or carbon dioxide asphyxiation. The eyes were enucleated and pretreated with 0.12% proteinase K (ICN Biomedicals Inc., Ohio, USA) for 15 min at 37 °C in R16, a serum-free culture medium (Gibco, Paisley, UK). Proteinase K activity was blocked by adding 10% fetal bovine serum, and the eyes were rinsed in a serum-free medium. After cornea, lens, sclera, and choroid dissection, the retina with the RPE attached was transferred onto a culture membrane insert (Corning Life Sciences, Lowell, USA) with the RPE facing the membrane. The membrane inserts were placed into six-well culture plates with R16 medium (Gibco) and incubated at 37 °C in a humidified 5% CO_2_ incubator. The culture medium was changed every 2 days during the 10 culturing days. Retinal explants were treated with 10 µM GSK-J4 (≥98% (HPLC); Sigma). GSK-J4 was dissolved in 0.2% dimethyl sulfoxide (DMSO; Sigma) and diluted in an R16 culture medium. The exact amount of DMSO was diluted in the culture medium for control explants. Explants were collected at PN24, fixed with 4% paraformaldehyde (PFA), and cryoprotected with graded solutions of 10, 20, and 30% sucrose before embedding in tissue freezing medium (Leica Microsystems Nussloch GmbH, Nussloch, Germany).

### 2.3 Intravitreal injections

Mice were anesthetized with an intraperitoneal injection of 40mg/kg Ketamine (Ceva Animal Health Pty Ltd, Amersham, UK) and 5mg/kg Ilium Xylazil-100 (Troy Laboratories, New South Wales, Australia) or subcutaneously with a mixture of ketamine (85mg/kg; Parke-Davis, Berlin, Germany) and xylazine (4mg/kg; Bayer AG, Leverkusen, Germany). Single intravitreal injections of 0.5uL were administered at PN14, with either 100nM of Trichostatin A (TSA; Sigma) or 10µM of GSK-J4 (Selleckchem, Texas, USA) administered in one eye and the contralateral eye receiving 0.0001% DMSO (diluted in 0.9% sodium chloride solution [Sigma]). GSK-J4 contained 0.0001% DMSO and was further diluted in 0.9% sodium chloride solution to make a 100µM solution, while TSA also contained 0.0001% DMSO with dilution to make a 100nM solution. Assuming a 5uL free vitreous volume [26], the final concentration of treatment was 10nM TSA or 10µM of GSK-J4. Anesthesia was reversed with Ilium Atipamezole (1mg/kg; Troy Laboratories) or Antisedan (Atipamezole; 2mg/kg; Orion Corp, Espoo, Finland).

### 2.4 Immunohistochemistry

Mice were euthanized via cervical dislocation or carbon dioxide asphyxiation at PN24, with eye orientation being marked before enucleation. Eyes were fixed in 4% PFA (Electron Microscopy Science Inc., Pennsylvania, USA) for 30 minutes before the cornea and iris were dissected. Eyes were fixed for another 30 minutes in 4% PFA before being placed into a sucrose gradient (Sigma) for cryoprotection. After 24 hours, the lens was removed, and the posterior eyecup was frozen in Tissue-Tek® O.C.T™ Compound (Sakura Finetek, California, USA) before being cryosectioned on the sagittal plane. Retinal sections were rehydrated with 1X phosphate buffered saline (PBS) for 20 minutes at room temperature (RT) before blocking solution was applied for 1 hour at RT. Blocking solution contained 0.5% Triton X 100 (ThermoFisher, Massachusetts, USA), 1% Bovine Serum Albumin (Bovogen Biologicals Pty Ltd., Victoria, Australia), and 10% Normal Goat Serum (Sigma), diluted in 1X PBS. Primary and secondary antibodies were applied to detect antigens in samples (complete list of antibodies in Table S1). Following application of primary and secondary antibodies, slides were incubated for 5 minutes with DAPI (4’, 6-diamidino-2-phenylindole, 0.5ug/mL in 1X PBS) and were mounted using fluorescence mounting medium (Dako). Negative controls were included in each batch by omitting the primary antibody.

### 2.5 Microscopy and Cell Counting

Images from retinal explants were captured using Z-stacks on a Zeiss Axio Imager Z1 ApoTome Microscope using the 20X air objective. The number of cones was quantified on 6-16 different positions, from 3-6 retinal cross-sections obtained from different positions within a retinal explant. *In vivo* images were taken on the Nikon Ti-E inverted motorized microscopy with Nikon A1Si spectral detector confocal system, with Z-stacks obtained on the 20X air objective. Quantification was performed as previously described [18], where each biological replicate had two sections imaged, and each section had an image taken in four positions, relative to the optic nerve: +10° (superior, central), +80° (superior, peripheral), −10° (inferior, central) and −80° (inferior, peripheral) positioning. The operator was blinded, and the numbers of GFP-positive cells (cones) were counted manually. The length of the outer nuclear layer (ONL) was manually measured using NIS-C Elements software. Cone quantification data were expressed as the number of cones/mm. Calculation of H3K9 and H3K27me3 positive cones was performed by counting a population of cones and determining the percentage of positive cells. All qualitative histology presented is representative images of staining throughout the tissue and from at least three biological replicates.

### 2.6 Electroretinography (ERG)

Retinal function in uninjected, sham-injected, and GSK-J4 injected mice was assessed at PN24 in the *Pde6c*.GFP line. Both full-field flash scotopic and photopic recordings were performed on the Celeris full-field ERG system (Diagnosysis LLC, Massachusetts, USA). Mice were dark-adapted overnight, and handled under dim red light for scotopic readings. Mice were anesthetized with an intraperitoneal injection of 80mg/kg Ketamine (Ceva Animal Health Pty Ltd) and 10mg/kg Ilium Xylazil-100 (Troy Laboratories), and pupils were dilated by application of a drop of 1% tropicamide to the cornea (Alcon, Western Australia). A drop of 2% hypromellose (HUB pharmaceuticals LLC, California, USA) was also applied to the cornea and eye electrodes to ensure moisture was retained and to act as a contact fluid. Animals were kept warm throughout the procedure with the in-built platform heater on the Celeris system. For scotopic readings, single-flash intensities were obtained through 1ms flashes with intensities of 0.01, 0.1, 0.3, 1, 3, 10, and 25 cd.s.m^-2^. The time between consecutive flashes was 10 seconds, while the stimulus was repeated four times at 0.10Hz. 60 seconds recovery time between different flash intensities was allowed. Following scotopic readings, mice were light-adapted for 10 minutes at 30 cd.s.m^-2^. A series of flashes on a background of 30 cd.m^-2^ were performed at 2 Hz and at intensities of 3 and 10 cd.s.m^-2^. The time between consecutive flashes was 0.5 seconds, while the stimulus was repeated 32 times. Flicker ERG responses were recorded at a pulse frequency of 10 and 30Hz, with a background of 30cd.m^-2^ and a pulse intensity of 3cd.s.m^-2^. Analysis of a- and b-waves was performed on the Espion V6 software (Diagnosys LLC).

### 2.7 Cell dissociation and single-cell sorting of cones

Fresh retinae were dissected before being placed in activated papain/DNase (1:20 solution; Worthington Biochemicals, New Jersey, USA) and incubated at 37°C for 45 minutes on a shaker. After incubation, the solution was gently triturated before spinning at 1500rpm for 5 minutes. The supernatant was discarded before the cell pellet was resuspended in Earle’s balanced salt solution (EBSS; Worthington Biochemicals) that contained 9.5%v/v Ovomucoid Inhibitor and 5%v/v DNase (Worthington Biochemicals). This resuspended solution was incubated at 37°C for 10 minutes and then re-spun at 1500rpm for 5 minutes. The supernatant was discarded, and the cell pellet was resuspended in EBSS and 10%v/v DNase. Samples were incubated with eBioscience™ Fixable Viability Dye eFluor™ 660 (Invitrogen, California, USA) on ice and in the dark for 30 minutes. Samples were then washed in fluorescent activated cell sorting (FACS) buffer that contained 2% heat-inactivated fetal calf serum (Fisher Biotec, Wembley, Western Australia), 1mM EDTA (Invitrogen), and 1X PBS. Cells were pelleted at 1400xrpm for 5 minutes, the supernatant discarded and resuspended in FACS buffer. Cell sorting was performed on a BD FACSMelody Cell Sorter, with single GFP-positive cells collected into 96-well plates containing 3uL of lysis buffer and ribonuclease inhibitor (one cell/well). Sorted plates were stored at −80°C until processed.

### 2.8 Single-cell RNA sequencing and bioinformatics

Sample preparation (RNA extraction, library preparation) and sequencing was performed at the Institute for Immunology and Infectious Diseases (Murdoch University, Perth, Western Australia) using the method described in Wanjalla et al. 2021 [27], which is an adapted version of the SMARTseq2 and MARS-seq approaches [28, 29]. Briefly, the assay utilized uniquely tagged primers for reverse transcription and template switching with a pre-amplification step to increase the yield and transcript length of the single-cell cDNA library. During the initial reverse transcription step, cDNA was tagged with well-specific barcodes coupled with a unique molecular identifier (UMI) to allow for multiplexing and increased sample throughput. Samples were then pooled and amplified using the KAPA HiFi HotStart ReadyMix (Roche, Basel, Switzerland), as per the manufacturer’s instructions. UMIs enabled quantitation of individual gene expression levels within single cells, thereby reducing technical variability and bias introduced during the amplification step [30-32]. Purified amplicons from the 5’ and 3’ were pooled to equimolar amounts, and indexed libraries were created for sequencing using Truseq adapters and quantified using the Kapa universal qPCR library quantification kit (Kapa Biosystems Inc., MA, USA). Samples were sequenced on an Illumina Novaseq using a 2×100 paired-end chemistry kit (Illumina Inc., CA, USA), as per the manufacturer’s instructions. Reads were quality-filtered and passed through a demultiplexing tool to assign reads to individual wells and to the 3’end and 5’ end. Reads for the individual single-cells were demultiplexed using plate-ID (30 nt), and cell barcode (6 nt). The reads were further demultiplexed as either 3’ or 5’ using primer sequence (30 nt), and the reminder 45 nt sequences were aligned to the GRCh38 Mus-Musculus reference genome (Ensembl rel. 97) using the CLC Genomics Workbench (CLC Bio) (v.20, QIAGEN Bioinformatics). An eight-nucleotide UMI tag and mapping coordinates were used to remove PCR-duplicate reads. Gene-specific read counts were calculated using RSubread:featureCount using Gencode M17 (2018) annotations, and the 3’ and 5’ counts were summed. Genes with > 0 counts in fewer than three cells and cells that either contained less than 200 genes or more than 5% mitochondrial content were filtered out. Downstream analyses (normalization, principal component analysis, differential expression, and visualization) were performed in Seurat v.2.3.4 R package [33]. A significance threshold of unadjusted *P*<0.01 and absolute log fold >0.3 was used to identify differentially expressed genes (DEGs), with batch correction applied using the Combat approach [34]. Gene ontology analysis was performed using Enrichr [35], with redundant terms trimmed using Revigo [36] (unadjusted *P*<0.01). Ingenuity Pathway Analysis (Qiagen) was used for canonical pathway and network analysis. To identify and define any subpopulations, cells were clustered on the basis of their expression profiles using Seurat to perform unsupervised UMAP analysis, along with Clustering Trees software to identify the optimal number of clusters [37]. The Seurat FindAllMarkers function was used to identify gene expression markers that were more highly expressed in each cluster, relative to all other clusters, indicating transcriptionally related cells regardless of genotype (a marker gene defined as being >0.3 log-fold higher (unadjusted *P*<0.01) than the mean expression value in the other sub-clusters, and with a detectable expression in > 25% of all cells from the corresponding sub-cluster). All raw data has been deposited into the Gene Expression Omnibus (GEO) repository GSE195895.

### 2.9 Statistical Analysis

Statistical analysis was performed using the PRISM software (GraphPad; San Diego, USA) and Excel (Microsoft; Washington, USA). Either t-test or ANOVA was used to analyze result significance (*P*<0.05, except for single-cell RNA sequencing where *P*<0.01).

## 3. Results

### 3.1 HDAC inhibition increases H3K27 trimethylation in Pde6c^cpfl1^ cones

Interaction between complex and diverse epigenetic modifications results in active transcription (via the activity of histone acetyltransferases, lysine methylases, lysine demethylases, and ubiquitin) or repressed transcription (histone deacetylases, lysine methylases, lysine demethylases, and de-ubiquitinating enzymes; Fig. 1A). To study this relationship specifically in cone photoreceptor degeneration, we have investigated the effect of *in vivo* delivery of the pan-HDAC inhibitor Trichostatin A (TSA) on the methylation status on histone H3 lysine residues in *Pde6c*^*cpfl1*^ mice. A single intravitreal injection of 10nM TSA at PN14 led to a significant increase in cone survival up to PN30 compared to sham treatment (Fig. 1B, C; One-way ANOVA **<0.01, n= 4). This finding additionally emphasizes the beneficial effect of HDAC inhibition on photoreceptor cell survival in multiple models of IRD [15-17]. We assessed the levels of H3K27 methylation in wt and *Pde6c*^*cpfl*^ controls (uninjected and sham-injected) and TSA-treated cones by co-staining with H3K27me3-specific and cone-specific glycogen phosphorylase (Glyphos) antibodies [16, 38] (Fig. 1B). H3K27 trimethylation was detected in the cones of wt mice, while in uninjected and sham-injected *Pde6c*^*cpfl1*^ mice, staining of H3K27me3 was seldomly observed in the cones. Treatment with 10nM TSA resulted in a partial restoration of H3K27me3 levels (Fig. 1D) with 44% of the TSA-treated cones expressing H3K27me3 in comparison to 25% H3K27me3 positive cones in sham-injected retinae (Fig. 1E; Welch’s T-test, *<0.05, n=3). We evaluated another methylation site, H3K9me3, with similar gene silencing properties as H3K27me3, and observed a trend towards more H3K9me3 positive cones after the treatment (Fig. S1). As the H3K27 trimethylation status in the cones of *Pde6c*^*cpfl1*^ mice appeared to be markedly downregulated, we further investigated if administration of a drug that inhibits demethylation at H3K27 sites would be a suitable neuroprotective treatment to prevent cone loss.

**Fig. 1.**
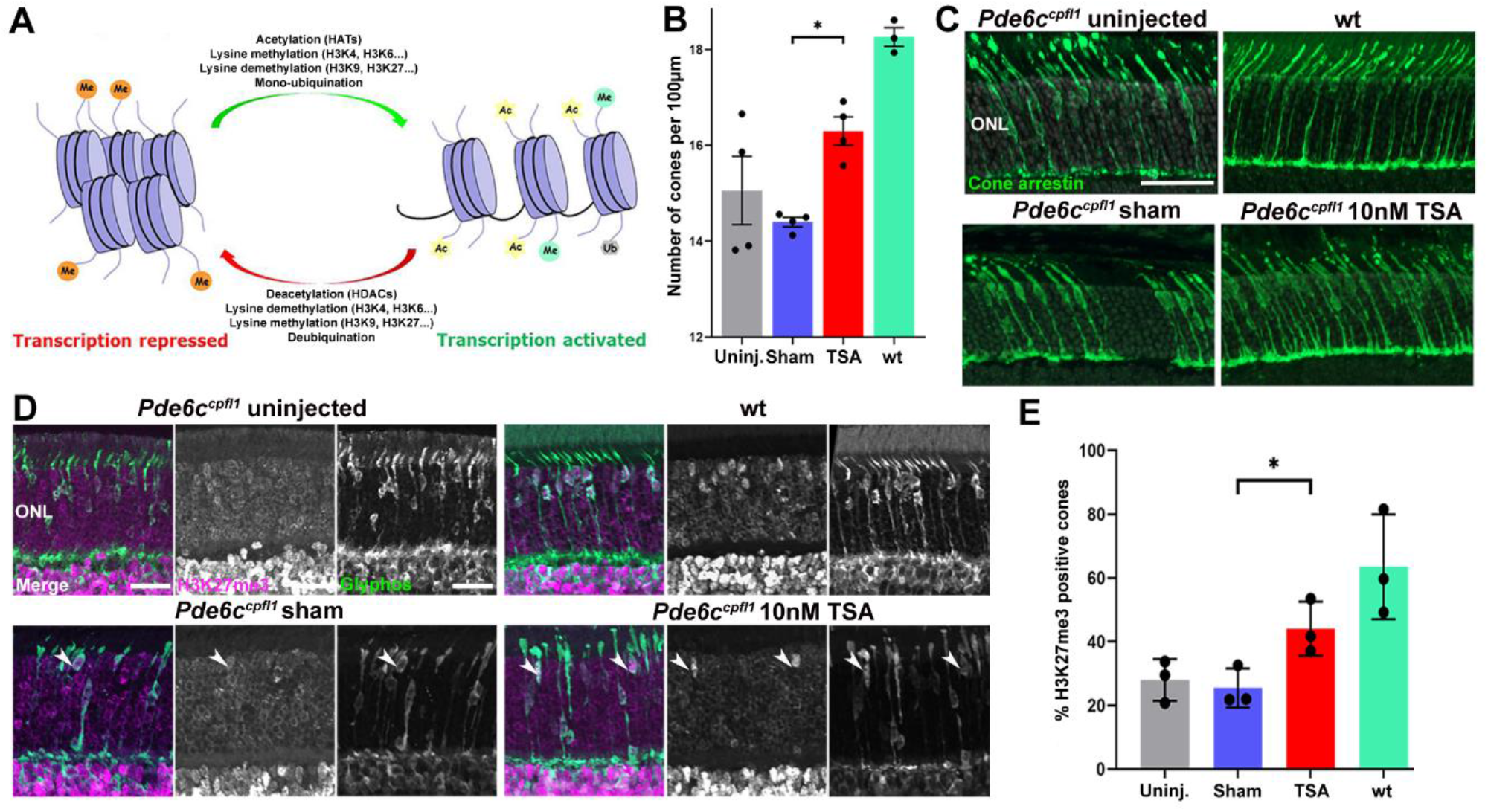
In vivo intravitreal delivery of Trichostatin A (TSA) at PN14 increases H3K27 trimethylation and cone numbers at PN24 in the *Pde6c*^*cpfl1*^ mouse model of cone dystrophy. **A** The balance and interplay between repressed and activated transcription. **B, C** A single intravitreal injection of Trichostatin A (TSA) led to significant cone (green) retention in *Pde6c*^*cpfl1*^ mice compared to sham-injected controls. One-way ANOVA, n=4, \**P*<0.01. Scale bar 50µm. ONL, outer nuclear layer. **D** Uninjected and sham-injected *Pde6c*^*cpfl1*^ mice showed scarce H3K27me3 methylation (magenta) costaining with Glyphos (glycogen phosphorylase in green). However, wt and TSA-treated cones displayed rich H3K27 tri-methylation. Arrows indicate cells that show co-localization of H3K27me3 and Glyphos. Scale bar 20µm. **E** Quantification of the percentage of H3K27me3 positive cones showed a significant increase in H3K27me3 localization in the cone photoreceptors, in TSA-treated retinae compared to sham-injected controls. Welch’s T-test, n=3, \**P*<0.01

### 3.2 The histone demethylase inhibitor GSK-J4 induces global transcriptional changes in the Pde6c^cpfl1^ cones

We have previously shown that inhibition of histone deacetylases in a mouse model of RP results in transcriptional changes of numerous cone-specific genes [18]. To assess to which extent direct modulation of histone methylation is associated with changes in cone-specific transcription profiles, we performed a single intravitreal injection in the *Pde6c*^*cpfl1*^ mice with the Jumonji H3K27me2/me3 demethylase inhibitor GSK-J4 at the onset of cone degeneration, PN14 (Fig. 2A) [21]. At PN24, the peak of cone death, we tested cone function and gene expression to assess the effects of the treatment (Fig. 2A). We used our previously validated transgenic *Pde6c*.GFP mouse, which is phenotypically the same as the *Pde6c*^*cpfl1*^ mouse. Our *Pde6c*.GFP mouse carries the same mutations in the *Pde6c* gene as the *Pde6c*^*cpfl1*^ alongside a copy of the e*GFP* gene downstream of the promoter for the *Chrnb4* gene, driving GFP expression exclusively in the cones. This allows for easy quantification and cell sorting of cone photoreceptors [24]. Using GFP fluorescence, we employed fluorescent-activated cell sort (FACS) of GSK-J4 treated and uninjected retinae to isolate cones for single-cell RNA sequencing. After processing and quality control, a total of 85 untreated cones and 71 GSK-J4 treated cones were analyzed. With a significance of unadjusted *P*<0.01 and an absolute log fold change greater than 0.3, we identified 2269 differentially expressed genes (DEGs), with more than 75% of them being downregulated (Fig. 2B, C; full list in Table S2).

**Fig. 2.**
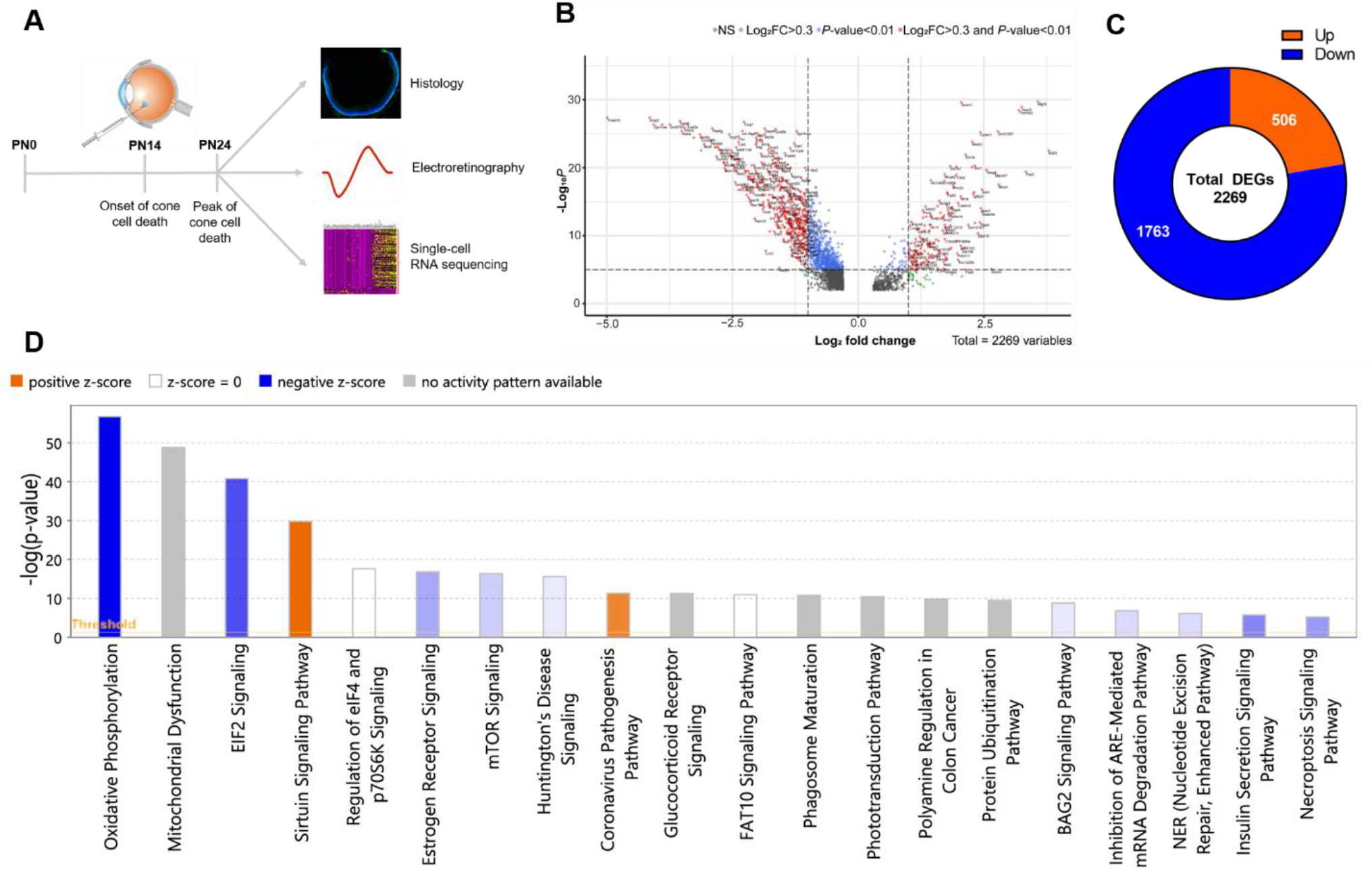
Administration of a single intravitreal injection of GSK-J4 at the onset of cone cell death in *Pde6c*.GFP mice induced global transcriptional changes. **A** Schematic overview of the experimental design of *in vivo* GSK-J4 intravitreal delivery at the onset of cell death (PN14) and sample collection at the peak of cell death (PN24). Eyes were collected for histological analysis, electroretinography, and single-cell RNA sequencing of the cones. **B** Volcano plot of differentially expressed genes after GSK-J4 treatment, based on their –Log_10_ *P-value* and Log_2_ fold change. Genes in red were analyzed further, as they were considered biologically relevant, using benchmarks of a *P-value* <0.01 and a fold change greater than 0.3 or less than −0.3. **C** Pie chart showing the number of genes that were upregulated (<25%) and downregulated (>75%) in our dataset. **D** Ingenuity Pathway Analysis bar graph, depicting the most enriched canonical pathways following the GSK-J4 administration (i.e., mitochondrial function and dysfunction, endoplasmic reticulum stress, epigenetic pathways, amongst others). The Z score represents the overall activation (yellow) or repression (blue) of each pathway, dependent on the direction of gene expression within the pathway. Intravitreal injection image available under license: Creative Commons Attribution-NonCommerical 3.0 Unported at https://creativecommons.org/licenses/by-nc/3.0/; https://researchgate.net/figure/Intravitreal-injection-Note-Intravitreal-injection-of-drug-is-recently-used-for_fig2_235729464

We evaluated upregulated and downregulated genes after the GSK-J4 treatment and utilized the Enrichr gene ontology (GO) database to determine the functional association of these genes. The most upregulated DEGs included genes associated with DNA damage repair (*Mgmt, Rpa2, Fem1b, Brip1, Ercc4, Taok1, Hinfp, Ube2n*), cell cycle (*Fem1b, Rpa2, Cdc23, Hecw2*), and T cell response (*Il23a, Il12rb1*). On the other hand, we observed a significant downregulation in various mitochondrial processes, including ones influenced by ATP synthase presence (*Atp5a1, Atp5b, Atp5c1, Atp5d, Atp5e, Atp5f1, Atp5g1, Atp5g2, Atp5g3, Atp5h, Atp5j, Atp5j2, Atp5k, Atp5l, Atp5o1, mt-Atp6, mt-Atp8*), cytochrome C function (*Cox4i1, Cox5a, Cox5b, Cox6a1, Cox6b1, Cox6b2, Cox6c, Cox7a2, Cox7a2l, Cox7b, Cox7c, Cox8a, Cox14, Cox19*), RNA polymerase II subunits (*Polr2c, Polr2d, Polr2e, Polr2f, Polr2g, Polr2h, Polr2j, Polr2k, Polr2l, Polr2m*) and other protein targeting and translational genes (full list of genes and GO in Table S2-S4). We also observed GSK-J4 induced expression changes in several histone modifiers, including histone (*Kmt1d, Kmt2d, Kmt5c, Prmt2*) and non-histone (*Eef1akmt*) methyltransferases, demethylases (*Kdm7c, Prdm11*), acetyltransferases (*Atf2, Atf4*), and deacetylases (*Hdac4, Sirt2*). All were downregulated with the exception of *Prdm11* and *Hdac4*. Two components of the MLL4-COMPASS complex (*Dpy30, Ash2l*) were also downregulated, including the H3K27me2/3 demethylase *Kdm6a* and H3K4me1/2/3 methyltransferase *Kmt2d* (Table 1 and Table S2).

**Table 1.** Histone modifying genes affected by the GSK-J4 treatment. The single-cell RNA sequencing data suggests that all genes displayed were downregulated, except *Hdac4* and *Prdm11*, which were upregulated (bolded)

| Histone modification | Gene symbol |
| --- | --- |
| Methyltransferases (Kmt) | *Kmt1d*, *Kmt2d*, *Kmt5c*, *Eef1akmt* |
| Demethylases (Kdm) | *Kdm7c* |
| Acetyltransferases (Hat) | *Atf2*, *Atf4* |
| Deacetylases (Hdac) | *Sirt2*, ***Hdac4*** |
| Other | *Ashl2*, *Dpy30*, *Prmt2*, ***Prdm11*** |

Canonical pathway analysis was performed using the Ingenuity Pathway Analysis tool (IPA; Qiagen), with the top 20 enriched pathways shown in Fig. 2D. More specifically, within the top three pathways, GSK-J4 treated cones showed downregulation of the oxidative phosphorylation, mitochondrial dysfunction, and EIF2 signaling pathways (Fig. S2 and S3; Full list of canonical pathways in Table S5). Further pathway analysis highlighted mitochondrial function, organization, and protein translation as key biological processes associated with GSK-J4 treatment (Fig. 3A). Downstream gene targets of GSK-J4 included translocases of mitochondrial membranes (*Timm13, Timm8b, Timm22, Timm50, Pam16, Tomm40, Tomm22*), NADH dehydrogenases (*mt-Nd1, mt-Nd2, mt-Nd3, mt-Nd4, mt-Nd5, mt-Nd6*), NADH-ubiquinone oxioreductases (*Ndufaf2, Ndufaf3, Ndufa7, Ndufb7, Ndufa11*), cytochrome related molecules (*mt-Co1, mt-Co2, mt-Co3, mt-Cyb, Uqcr11, Uqcrq*), amongst others (Fig. 3B).

**Fig. 3.**
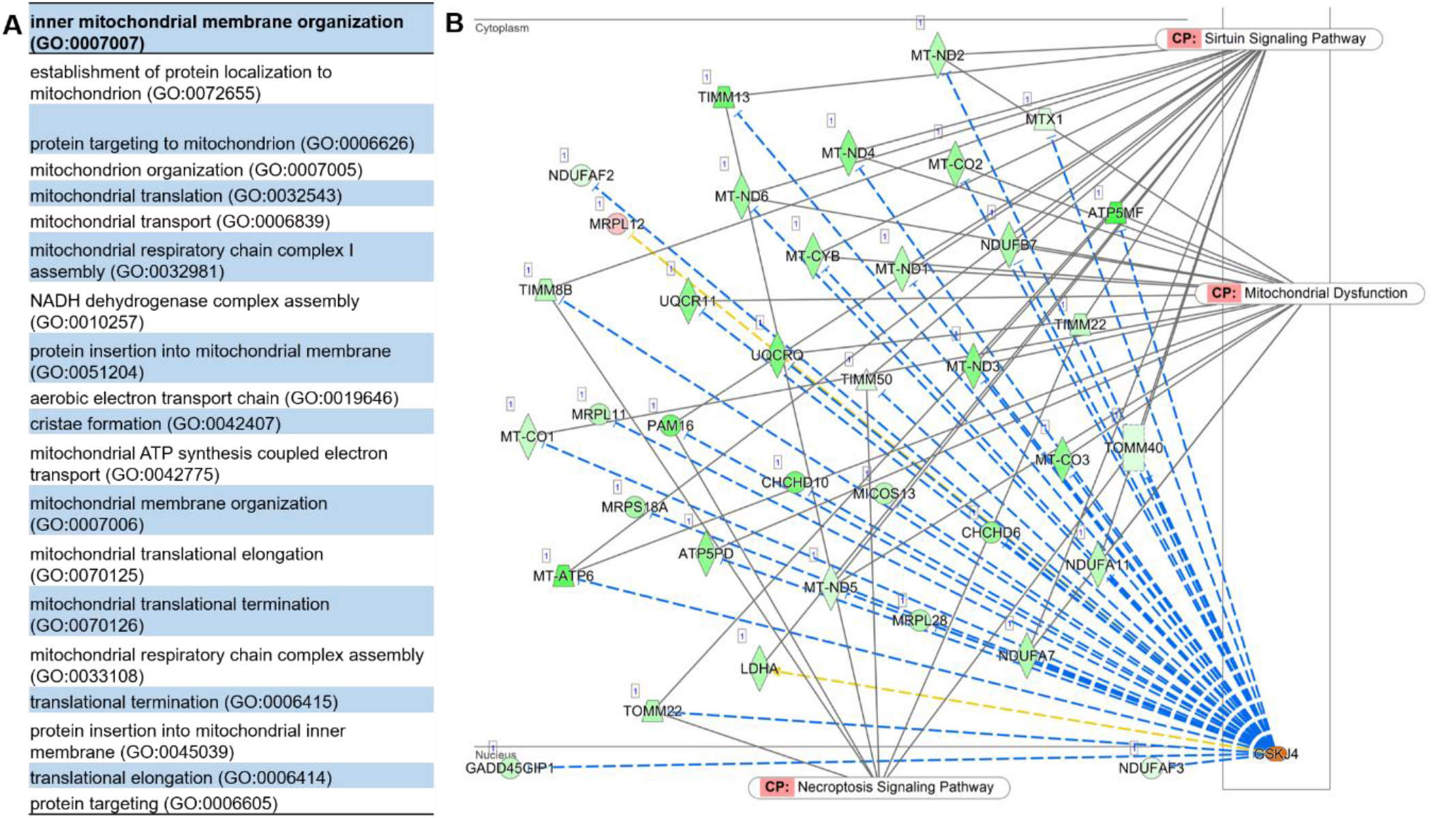
A complex interaction between different pathways following the GSK-J4 treatment. **A** The most enriched biological processes after GSK-J4 treatment, showing substantial enrichment of mitochondrial function and organization, and translation. Gene ontology analysis was performed using Enrichr, with redundant terms trimmed using Revigo (unadjusted *P<*0.01). **B** Network visualization of the downstream targets of GSK-J4 with the canonical pathways overlaid. Many of these genes are involved in the mitochondrial dysfunction pathway, while others show involvement in sirtuin and necroptosis signalling pathways. Schematic downloaded from QIAGEN Ingenuity Pathway Analysis. Green downregulated; red upregulated. Blue leads to inhibition, and yellow indicates inconsistent findings into a downstream state of a molecule

### 3.3 Gene expression clustering and downstream targets following GSK-J4 treatment

The single-cell approach allowed us to examine the cellular heterogeneity of cones in response to GSK-J4 treatment and to identify genetic markers that may be used to discriminate cone subpopulations. An unsupervised uniform manifold approximation and projection (UMAP) plot was used to visualize the Seurat clustering analysis, which plots transcriptionally similar cells using a weighted k-nearest neighbour method [39] (Fig. 4A). Clusters were identified based on their gene expression profile (top 50 genes for each cluster are shown in Table S6), defined as being >0.3 log-fold higher than the mean expression value in the other sub-clusters, and with a detectable expression in >25% of all cells from the corresponding sub-cluster. Annotation of the clusters revealed that cluster 0 predominantly contained GSK-J4 treated cones, whilst clusters 1 and 2 comprised primarily of untreated cones (Fig. 4B). Interestingly, untreated cone cells formed two distinct clusters whilst GSK-J4 treated cones formed one. Genes from the cluster heat map, representing the top 50 genes from each cluster (selected by average log fold change; Fig. 4C), were used for GO analysis to identify characteristics of each cluster (see Table S6 for the complete list of genes and GO). Biological processes associated with cluster 0 focused on DNA repair and damage response and positive regulation of apoptosis. Genes involved with oxidative phosphorylation (complexes I and IV of the aerobic electron transport chain) and activation of endoplasmic reticulum stress response defined cluster 1. Cluster 2 was enriched for respiratory chain function (complexes I and III) and negative regulation of apoptosis. Interestingly, the expression of other known histone modifiers, long non-coding RNA (lncRNA), was also a descriptor of each cluster, with cluster 0 containing six lncRNAs in the top 50 genes (*Gad1os, Gm5103, 4632411P08Rik, Gm31557, Gm47152, 4930528D03Rik*), compared to two lncRNAs for cluster 1 (*Gm29152, Gm15417*) and one lncRNA for cluster 2 (*Airn*).

**Fig. 4.**
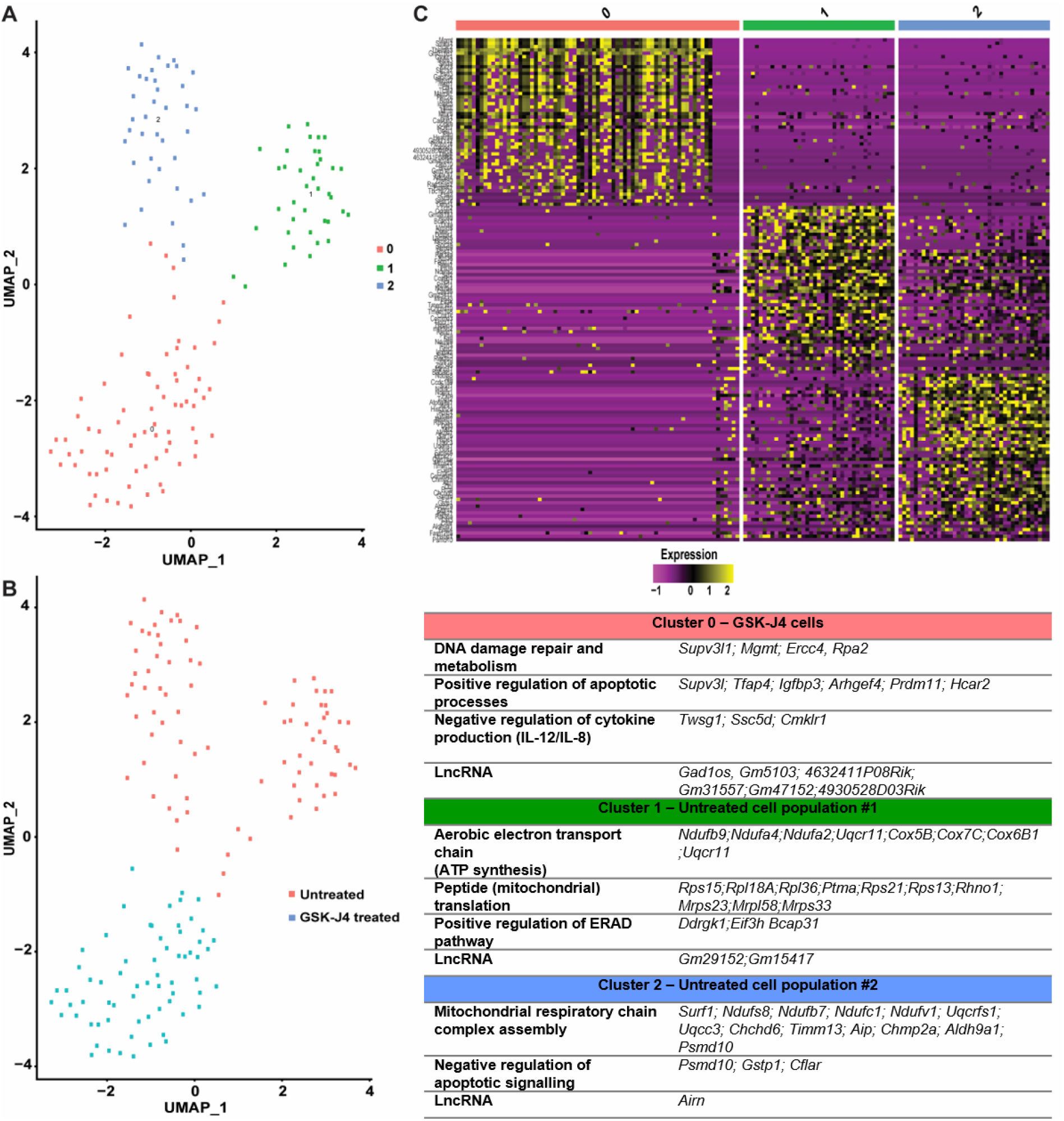
The GSK-J4 treatment-induced changes in cone photoreceptor clustering and gene expression. **A, B** Clustering analysis of cones visualized on a UMAP (uniform manifold approximation projection) plot revealed three clusters, with cluster 0 comprising the GSK-J4 treated cones, while the cones from uninjected *Pde6c*.GFP mice showed two different cell clusters, clusters 1 and 2. **C** Heat map of the top 50 most expressed genes that characterize each cluster, with the table showing the enriched genes and gene ontology (GO) biological processes associated with each cluster. Gene expression is scaled and presented as Log2-transformed fold change

### 3.4 In vivo assessment of the histone demethylase inhibitor GSK-J4 on cone photoreceptor survival and function

While treatment with GSK-J4 in *Pde6c*.GFP mice induced robust changes in cone-specific transcription profiles, we did not observe a difference in cone numbers compared to controls (Fig. 5B; 2-way ANOVA, *P*>0.05, n=7). We validated this with IHC staining, with all groups showing a similar number of cone cells, cone morphology, opsin localization, and glial fibrillary acidic protein expression as a marker for Müller glia activation (Fig. 5A, E). We also performed photopic (cone-function) and scotopic (rod-function) ERG recordings to assess any functional changes to the retina after treatment with GSK-J4. We did not observe a significant difference at any stimulus intensity in photopic or scotopic ERG between groups in either a-or b-wave amplitude, suggesting the treatment did not have any deleterious effects on photoreceptor function (Fig. 5C, D; photopic data not shown; 2-way ANOVA, *P*>0.05, n=4).

**Fig. 5.**
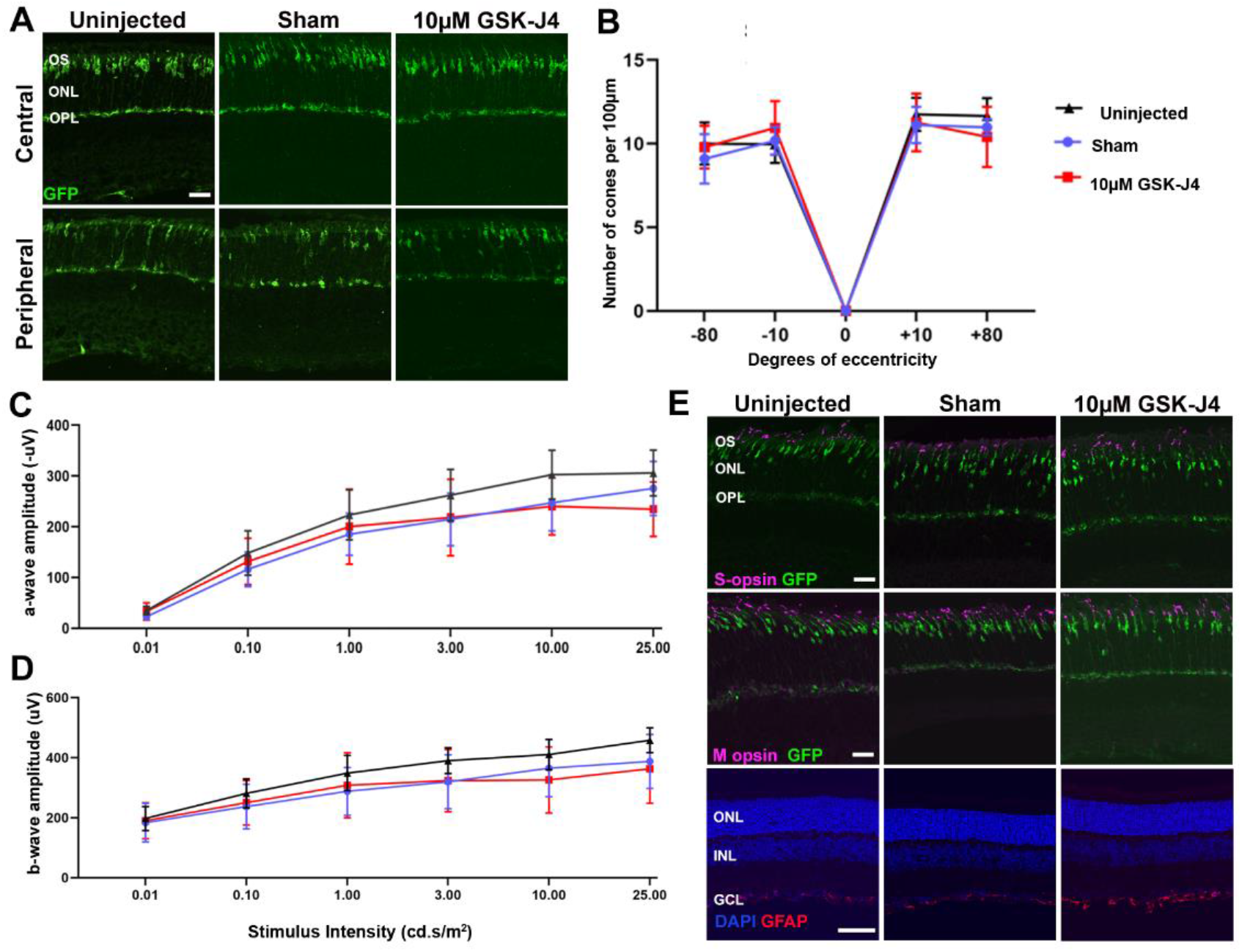
An intravitreal injection of GSK-J4 in *Pde6c*.GFP mice at PN14 does not affect morphology, cone numbers, protein localization or retinal function at PN24. **A** Representative central and peripheral retinal images from the *Pde6c*.GFP mouse line with GFP-positive cones (green) showing general morphology is similar between uninjected, sham-injected and GSK-J4 treated mice. Scale bar 20µm. OS, outer segment. ONL, outer nuclear layer. OPL, outer plexiform layer. GFP, green fluorescent protein. **B** Quantification of cone numbers per 100µm ONL length across the retina indicates that cone numbers do not differ between *Pde6c*.GFP uninjected, sham-injected or GSK-J4 treated retinae. Locations of the retina imaged were central superior (corresponding to +10° eccentricity from the optic nerve labelled with 0), peripheral superior (+80°), central inferior (−10°) and peripheral inferior (−80°). Two-way ANOVA, n=7, *P*>0.05. **C** Dark-adapted (scotopic) amplitude of the a-wave at different stimulus intensities, revealed no difference between uninjected, sham and GSK-J4 treated *Pde6c*.GFP mice. Two-way ANOVA, n=4, *P*>0.05. **D** Amplitude of the scotopic b-wave at six different stimulus intensities, showing no significant difference after treatment with GSK-J4. Two-way ANOVA, n=4, *P*>0.05. **E** S- and M-opsin (magenta) localization appear consistent between treated and uninjected groups. GFAP expression (red) is similar between uninjected and sham-injected mice, with a mild increase in expression after treatment with GSK-J4. Nuclei are stained with DAPI (blue). Scale bars 20µm. OS, outer segment. ONL, outer nuclear layer. OPL, outer plexiform layer. INL, inner nuclear layer. GCL, ganglion cell layer. GFAP, glial fibrillary protein

### 3.5 Continuous GSK-J4 treatment led to cone protection ex vivo

Although a single dose of GSK-J4 *in vivo* did not result in an increased cone survival at PN24, our single-cell RNA sequencing data suggested a beneficial effect of the drug via downregulation of disease-related pathways, including endoplasmic reticulum stress and mitochondrial dysfunction pathways. A single intravitreal injection may not have been sufficient to affect cone survival ten days post-injection due to the short half-life of GSK-J4. We assumed GSK-J4 followed extensive intravitreal clearance, shown for similar small lipophilic molecules [16, 40]. Hence, we tested the effect of continuous GSK-J4 treatment on cone survival. For this, we used *ex vivo* retinal explants of PN14 *Pde6c*^*cpfl1*^ mice, allowing the continuous presence of GSK-J4 treatment for ten days (Fig. 6A). Quantification on cones in GSK-J4 treated explants showed a 31.9% increase in cone survival when compared to sham controls (Fig. 6B, C; Unpaired T-test with Welch’s correction, ** *P*<0.01, n=11 sham, n=12 treated). We also observed an improved localization of M- and S-opsin and increased H3K27me3 staining in the cones after the treatment (Fig. 6D, E).

**Fig. 6.**
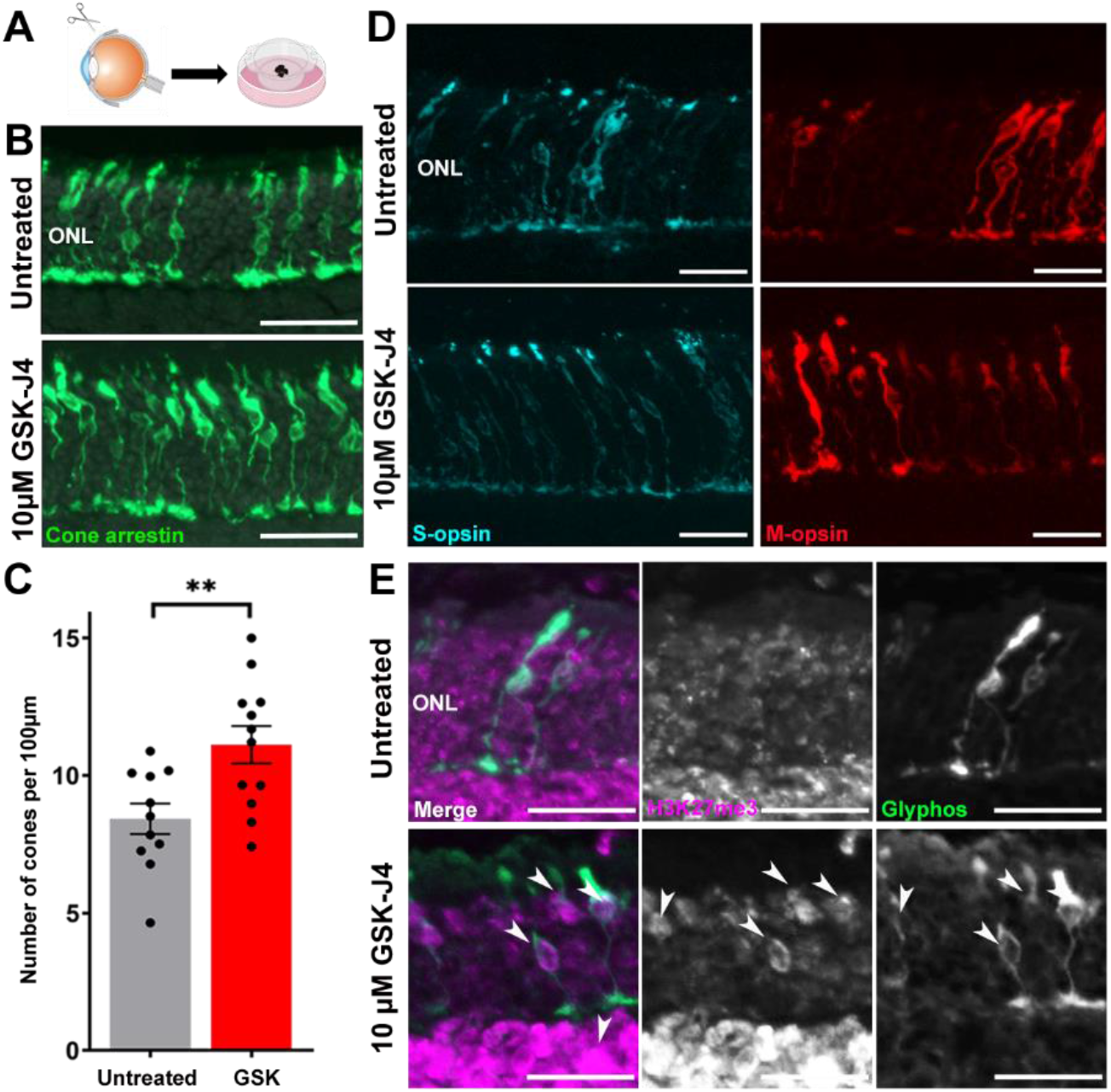
Continuous delivery of GSK-J4 to *Pde6c*^*cpfl1*^ retinal explants for ten days allowed significant retention of cone photoreceptors, improved morphology, and localization of M- and S-opsins. **A** Schematic diagram showing the process of dissecting the retina from PN14 *Pde6c*^*cpfl1*^ mice before growing retinal explants on culture membranes. Retinal explants were either sham-treated or treated with 10µM of GSK-J4 daily for ten days before collection at PN24. **B, C** Representative cone arrestin (green) immunostaining and quantification of cone numbers in *Pde6c*^*cpfl1*^ retinal explants show significant protection of GSK-J4 treated cones compared to untreated explants. Welch’s T-test, n=11 untreated, n=12 treated, \*\**P*<0.01. Scale bar 50µm. ONL, outer nuclear layer. **D** M-opsin (red) and S-opsin (cyan) immunostaining showed an improvement of protein localization in the cones after treatment with GSK-J4. Scale bar 20µm. ONL, outer nuclear layer. **E** An increase of H3K27me3 (magenta) staining was observed in *Pde6c*^*cpfl1*^ explants that had received GSK-J4 treatment (indicated by arrows) but not in untreated explants. Scale bar 20µm. ONL, outer nuclear layer. Glyphos, glycogen phosphorylase

## 4. Discussion

### Pde6c^cpfl1^ cones have reduced expression of H3K27me3, which is partially restored by TSA administration

Methylation and other key epigenetic markers are important in photoreceptor development, differentiation, fate determination, and function [41, 42]. Mature rods have a hypomethylated DNA epigenome, with a closed chromatin state when compared to the cones, indicating an overall repressed genetic environment [42]. Specifically, rods contain high levels of the repressive H3K9me3 in the nucleus, while cones show the presence of the active chromatin marker H3K4me3 [43, 44]. These differences between chromatin condensation and methylation marks give rise to the distinct function of rods and cones in light perception [43]. Consequent changes in these patterns may lead to deleterious effects with genes being incorrectly repressed or expressed. Indeed, a previous study in the *rd1* model of RP showed upregulation of the repressive methylation site H3K27me3 in rods whereby inhibition of histone methylation delayed rod degeneration [19]. Similarly, inhibition of lysine demethylase 1, which specifically demethylates H3K4me1/2 and H3K9me1/2, blocks rod degeneration in *rd10* mice [20]. Contrary to rods, our study shows that wt cones have high levels of H3K27me3, which is diminished in degenerating *Pde6c* mutant cones. This finding suggests that while increased levels of H3K27me3 levels appear to be deleterious in rods, the absence of H3K27 tri-methylation may be associated with cone degeneration. As the chromatin in mouse rods is already hypercondensed, the upregulation of repressive methylation marks such as H3K27me3 push nuclei towards pyknosis and subsequent cell death [45], while the same disease mechanism may not happen in mouse cones due to their different natural chromatin condensation.

Aberrant histone methylation status has been discussed as a possible pathological feature in various neurodegenerative diseases, including Alzheimer’s, Huntington’s, amyotrophic lateral sclerosis, and IRD [19, 46]. A particular point of focus is HDAC inhibition, classically utilized in cancer treatment, which has recently shown a neuroprotective capacity in many diseases, including IRD [47]. Several studies show the complex interactions between many epigenetic pathways, including the fact that histone methylation is a secondary target of HDAC inhibitors [48]. Our study shows that treatment with the HDAC inhibitor TSA provides significant retention of cone numbers [16, 17], while also causing an increase in H3K27me3 in the cones of the *Pde6c*^*cpfl1*^ mutant mouse. Other groups have observed this phenomenon, with Halsall et al. 2015 showing in lymphoblastoid cells that treatment with other pan-HDAC inhibitors SAHA and valproic acid, resulted in an upregulation of H3K27me3 at transcription start sites which are thought to promote cell survival through adaptive mechanisms via minimization of protein hyperacetylation, slowing growth and re-balancing patterns of gene expression [49]. Although TSA provided an increase in H3K27me3, our approach for this study was to take one step further and ask if directly inhibiting H3K27me3 demethylation via GSK-J4 administration could be used to promote cone survival pathways.

The effect of GSK-J4 on epigenetic modifiers

GSK-J4 was initially developed as a selective inhibitor of the H3K27me2/3 demethylases *Kdm6a* and *Kdm6b* but was later found to inhibit multiple members of the Jumonji-C domain-containing family, including the Kdm5 subfamily, which is responsible for the demethylation of active H3K4me1/2/3 markers [50]. As the methylation state of both H3K4 and H3K27 can be protective in neurodegeneration [20, 51], it indicates a role for GSK-J4 in concomitantly inhibiting both *Kdm6* and *Kdm5* subfamilies and suggest its neuroprotective action may involve the activity of both repressive H3K27me and active H3K4me.

Whilst we did not directly identify specific demethylases, our data points to the involvement of *Kdm6a*, both as a means of regulating H3K27me3 deposition and as a critical part of a protein group that regulates H3K4 methylation. In the MLL-COMPASS protein complex, KDM6A associates with the H3K4me1/2/3 methyltransferase, KMT2D, along with a core group of accessory proteins (WDR5, RbBP5, ASH2L, DPY30; The WRAD complex) [52]. Our single-cell RNA sequencing data showed that *Kmt2d*, along with two core protein genes (*Dpy30* and *Ash2l*), were downregulated after GSK-J4 treatment. This complex is involved in guiding RNA polymerase II to engage transcription [53], and we show downregulation of ten out of twelve RNA polymerase II subunits, suggesting a possible mechanism by which overall transcription-associated processes are decreased.

### Treatment of GSK-J4 leads to significant alterations in pathways associated with cell survival and mitochondrial function

After treatment with GSK-J4, we saw a substantial shift in the gene expression profile of single cones when visualized to investigate their relationship with one another. Untreated *Pde6c*.GFP cells clustered into two distinct sub-populations, with vastly different associated biological processes. Cluster 1 (untreated group 1) showed upregulation of the endoplasmic-reticulum-associated protein degradation pathway, which suggests an increase in cell death via misfolded or non-functional proteins [54]. Misfolded protein accumulation has been observed in other forms of IRD, with a study in P23H Rho models showing that this accumulation is a part of the disease progression [55]. On the other hand, cluster 2 (untreated group 2) showed enrichment of the mitochondrial assembly chain and a decrease in apoptosis. The dichotomy of cone cell scattering and associated biological processes suggests that at least at PN24, *Pde6c* mutant cones are at two different stages of the disease. One population of cells shows increased levels of cell death via protective mechanisms, indicating these cells might be in advanced stages of degeneration, while the other population has a decrease in apoptosis, meaning they may be in the early stages of the disease.

Interestingly, after treatment with GSK-J4, we observed one uniform population of cell clustering. The associated biological processes after GSK-J4 administration included DNA repair and damage responses and, surprisingly, positive regulation of apoptosis. A possible explanation for the increase in apoptosis after GSK-J4 treatment is that cells that cannot be repaired by DNA repair mechanisms undergo apoptosis, while salvageable cells are repaired [56]. We also observed a decrease in pro-inflammatory cytokines involved in IRD pathogenesis [57]. While GSK-J4 may facilitate apoptosis of cells that have gone past the point of no return, at the same time, it may have beneficial effects to still viable cells by reducing inflammatory conditions in the retina and enabling cell repair.

Overall, *Pde6c* mutant cones showed significant changes in several pathways associated with epigenetic modification, cell survival, and mitochondrial function as a consequence of the GSK-J4 treatment, with the most enriched canonical pathways being oxidative phosphorylation, mitochondrial dysfunction and EIF2 signaling, all of which were downregulated. Abnormal mitochondria function and eye disease have been linked numerous times. Both inherited, and age-related retinal diseases show abnormal mitochondria function and the build-up of reactive oxygen species (as reviewed by Lefevere et al. 2017) [58]. We observed the downregulation of many mitochondrial genes and processes, including ones related to cytochrome *c*. Cytochrome *c* is integral to healthy mitochondrial processes; however, overactivation has been extensively linked to cell death via activation of caspases [59]. An intriguing study by Huang et al. 2004 showed that the mitochondria play an important role in light-induced retinal degeneration models, suggesting that downregulation of three proteins; ATP synthase subunit-6, cytochrome *c* oxidase-III, and NADH dehydrogenase-3, was neuroprotective [60]. We observed a downregulation of their corresponding genes, *mt-Atp6, mt-CO3*, and *mt-Nd3*, suggesting this pathway may be providing a neuroprotective effect after treatment. We also observed downregulation of superoxide dismutases (SODs), involved predominantly in the mitochondrial dysfunction pathway. The role of SODs in RP has been extensively studied, with previous studies showing induction of cell death by increased expression of SOD2 [61], as well as data showing downregulation of SOD1 in the *rd10* mouse model [62]. Following GSK-J4 treatment, genes involved in the CHOP cell death pathway (part of the EIF2 signaling pathway) appear to be significantly repressed. Activation of the CHOP pathway can lead to cell death via long-term endoplasmic reticulum stress, causing induction of pro-apoptotic genes and suppression of anti-apoptotic proteins [63]. Other models of IRD have indicated an involvement of this pathway in cellular degeneration, with the *rd16* model of Leber’s congenital amaurosis showing an increase in the components of the CHOP pathway, leading to increased cell death [64]. Previous studies into the mechanism of action of GSK-J4 in an acute myeloid leukemia model indicate that GSK-J4 induces cell cycle arrest and apoptosis in cancer cells via the CHOP pathway [65]. This concept is intriguing, in that regulation of this pathway in cancer cells induces cell death, while in our models of cone degeneration it may prevent neurodegeneration.

### Continuous GSK-J4 treatment has a significant effect on cone survival in Pde6c^cpfl1^ retinal explants

While the data obtained by single-cell RNA sequencing suggested protective effects of GSK-J4 via the regulation of aberrant methylation and demethylation and indicated its anti-apoptotic properties, we did not see a significant increase in cone survival ten days after a single intravitreal injection of GSK-J4. Due to the fact that short-term exposure to GSK-J4 treatment was able to affect key pathways potentially involved in the pathogenesis of IRD, we evaluated the effect of continuous GSK-J4 treatment by assessing cone survival in *ex vivo* retinal explants after ten days of treatment. The observed ∼32% increase in cone numbers compared to sham controls suggests that GSK-J4 treatment may be beneficial when administered regularly or through drug delivery systems that allow continuous administration. Because GSK-J4 is capable of inducing changes at multiple levels, it may represent a broader treatment option that could be used in multiple IRD forms or in complex cone-loss diseases like age-related macular degeneration. Further studies should be conducted to assess the effectiveness of GSK-J4 in other models of IRD while utilizing a continual drug delivery system. It also remains to be investigated if combining neuroprotective agents, such as GSK-J4, with gene therapies can provide improved recovery of visual function compared to gene therapy alone. While human trials of the first FDA approved retinal gene therapy, Luxturna, showed significant visual improvement, it was reported that retinal degeneration still occurs despite the treatment [66]. A combinatorial treatment strategy may therefore be a way forward in such situations, which would allow for the retention of cells and prevent the eventual degradation of the patient’s vision due to cellular loss [66].

## Abbreviations

IRD, cGMP, HDAC, HAT, HDM, HMT, H3K, RP, PN, TSA, wt, RPE, ONL, ERG, DEGs, GO, UMAP, SODs

## Acknowledgements

We are thankful to Norman Rieger for technical assistance and to Francois Paquet-Durand and Yvan Arsenijevic for scientific discussions. The authors acknowledge the facilities, and the scientific and technical assistance of Microscopy Australia at the Centre for Microscopy, Characterisation & Analysis, The University of Western Australia, a facility funded by the University, State, and Commonwealth Governments. Funding for this work was provided by the Australian Government University Postgraduate Award, a generous donation by the Tahija Foundation, grants from the Future Health Research and Innovation Fund scheme through WA Near-miss Award Program 2019 round, The Lindsay & Heather Payne Medical Research Charitable Foundation (IPAP2020/1082), The Kerstan Foundation, The ProRetina Foundation, Deutsche Forschungsgemeinschaft (DFG TR 1238/4-1), and Swiss National Science Foundation (31003A 173008).

## 6. Statements and Declarations

## Author’s contributions

The conception and design of the work were performed by Dragana Trifunović, Annie L. Miller, and Livia S. Carvalho. All other authors contributed in various ways to the acquisition of data, analysis, interpretation, funding, and critical revision. Annie L. Miller wrote the first draft of the manuscript with the assistance of Paula I. Fuller-Carter, and all authors have read and approved the submission of the current manuscript.

## Data availability

The datasets generated and analyzed during the current study are available in the Gene Expression Omnibus repository and will be accessible upon this paper being published in a journal article.

## Ethics approval

All work performed for this study complied with the governing body in each respective country. The Harry Perkins Institute of Medical Research ethics committee approved all animal work undertaken in Australia (approval code AE052 and AE135b). The mouse experimental procedure at the University of Zürich was approved by the Veterinary Office of the Canton of Zürich (approval no. 141/2016). All mouse work performed at The University of Tübingen was approved by the Tübingen University committee on animal protection (approval no. AK09/18M).

## Consent to participate

Not applicable.

## Consent for publication

Not applicable.

## References

[1] Berger W, Kloeckener-Gruissem B and Neidhardt J (2010) The molecular basis of human retinal and vitreoretinal diseases. Prog Retin Eye Res 29:335–75. 10.1016/j.preteyeres.2010.03.004.

[2] Galvin O, Chi G, Brady L, Hippert C, Del Valle Rubido M, Daly A and Michaelides M (2020) The Impact of Inherited Retinal Diseases in the Republic of Ireland (ROI) and the United Kingdom (UK) from a Cost-of-Illness Perspective. Clinical ophthalmology (Auckland, N.Z.) 14:707–719. 10.2147/OPTH.S241928.

[3] Scott AW, Bressler NM, Ffolkes S, Wittenborn JS and Jorkasky J (2016) Public Attitudes About Eye and Vision Health. JAMA Ophthalmology 134:1111–1118. 10.1001/jamaophthalmol.2016.2627.

[4] Farvardin M, Afarid M, Attarzadeh A, et al. (2018) The Argus-II Retinal Prosthesis Implantation; From the Global to Local Successful Experience. Front Neurosci 12:584. 10.3389/fnins.2018.00584.

[5] Georgiou M, Fujinami K and Michaelides M (2021) Inherited retinal diseases: Therapeutics, clinical trials and end points—A review. Clinical & Experimental Ophthalmology 49:270–288. https://doi.org/10.1111/ceo.13917.

[6] Chang G-Q, Hao Y and Wong F (1993) Apoptosis: Final common pathway of photoreceptor death in rd, rds, and mutant mice. Neuron 11:595–605. https://doi.org/10.1016/0896-6273(93)90072-Y.

[7] Marigo V (2007) Programmed Cell Death in Retinal Degeneration: Targeting Apoptosis in Photoreceptors as Potential Therapy for Retinal Degeneration. Cell Cycle 6:652–655. 10.4161/cc.6.6.4029.

[8] Arango-Gonzalez B, Trifunović D, Sahaboglu A, et al. (2014) Identification of a common non-apoptotic cell death mechanism in hereditary retinal degeneration. PLoS One 9:e112142. 10.1371/journal.pone.0112142.

[9] Sancho-Pelluz J, Alavi MV, Sahaboglu A, et al. (2010) Excessive HDAC activation is critical for neurodegeneration in the rd1 mouse. Cell Death Dis 1:e24. 10.1038/cddis.2010.4.

[10] Paquet-Durand F, Hauck SM, van Veen T, Ueffing M and Ekström P (2009) PKG activity causes photoreceptor cell death in two retinitis pigmentosa models. J Neurochem 108:796–810. 10.1111/j.1471-4159.2008.05822.x.

[11] Paquet-Durand F, Sanges D, McCall J, Silva J, van Veen T, Marigo V and Ekström P (2010) Photoreceptor rescue and toxicity induced by different calpain inhibitors. J Neurochem 115:930–40. 10.1111/j.1471-4159.2010.06983.x.

[12] Brunet AA, Harvey AR and Carvalho LS (2022) Primary and Secondary Cone Cell Death Mechanisms in Inherited Retinal Diseases and Potential Treatment Options. Int J Mol Sci 23:10.3390/ijms23020726.

[13] Haberland M, Montgomery RL and Olson EN (2009) The many roles of histone deacetylases in development and physiology: implications for disease and therapy. Nature Reviews Genetics 10:32–42. 10.1038/nrg2485.

[14] Brown B, Aaron M (2001) The politics of nature. In: Smith J (ed) The rise of modern genomics, 3rd edn. Wiley, New York, pp 230–257.

[15] Lee MG, Wynder C, Bochar DA, Hakimi MA, Cooch N and Shiekhattar R (2006) Functional interplay between histone demethylase and deacetylase enzymes. Mol Cell Biol 26:6395–402. 10.1128/mcb.00723-06.

[16] Trifunović D, Arango-Gonzalez B, Comitato A, et al. (2016) HDAC inhibition in the cpfl1 mouse protects degenerating cone photoreceptors in vivo. Hum Mol Genet 25:4462–4472. 10.1093/hmg/ddw275.

[17] Trifunović D, Petridou E, Comitato A, Marigo V, Ueffing M and Paquet-Durand F, J. D. Ash, R. E. Anderson, M. M. LaVail, C. Bowes Rickman, J. G. Hollyfield and C. Grimm (2018) Primary Rod and Cone Degeneration Is Prevented by HDAC Inhibition. Retinal Degenerative Diseases 367–373.

[18] Samardzija M, Corna A, Gomez-Sintes R, et al. (2020) HDAC inhibition ameliorates cone survival in retinitis pigmentosa mice. Cell Death & Differentiation 10.1038/s41418-020-00653-3.

[19] Zheng S, Xiao L, Liu Y, Wang Y, Cheng L, Zhang J, Yan N and Chen D (2018) DZNep inhibits H3K27me3 deposition and delays retinal degeneration in the rd1 mice. Cell death & disease 9:310–310. 10.1038/s41419-018-0349-8.

[20] Popova EY, Imamura Kawasawa Y, Zhang SS-M and Barnstable CJ (2021) Inhibition of Epigenetic Modifiers LSD1 and HDAC1 Blocks Rod Photoreceptor Death in Mouse Models of Retinitis Pigmentosa. The Journal of Neuroscience 41:6775. 10.1523/JNEUROSCI.3102-20.2021.

[21] Trifunović D, Dengler K, Michalakis S, Zrenner E, Wissinger B and Paquet-Durand F (2010) cGMP-dependent cone photoreceptor degeneration in the cpfl1 mouse retina. J Comp Neurol 518:3604–17. 10.1002/cne.22416.

[22] Chang B, Hawes NL, Hurd RE, Davisson MT, Nusinowitz S and Heckenlively JR (2002) Retinal degeneration mutants in the mouse. Vision Res 42:517–25. 10.1016/s0042-6989(01)00146-8.

[23] Chang B, Grau T, Dangel S, et al. (2009) A homologous genetic basis of the murine cpfl1 mutant and human achromatopsia linked to mutations in the PDE6C gene. Proc Natl Acad Sci U S A 106:19581–6. 10.1073/pnas.0907720106.

[24] Brunet AA, Fuller-Carter PI, Miller AL, Voigt V, Vasiliou S, Rashwan R, Hunt DM and Carvalho LS (2020) Validating Fluorescent Chrnb4.EGFP Mouse Models for the Study of Cone Photoreceptor Degeneration. Translational Vision Science & Technology 9:28–28. 10.1167/tvst.9.9.28.

[25] Siegert S, Scherf BG, Del Punta K, Didkovsky N, Heintz N and Roska B (2009) Genetic address book for retinal cell types. Nature Neuroscience 12:1197–1204. 10.1038/nn.2370.

[26] Kaplan HJ, Chiang C-W, Chen J and Song S-K (2010) Vitreous Volume of the Mouse Measured by Quantitative High-Resolution MRI. Investigative Ophthalmology & Visual Science 51:4414–4414.

[27] Wanjalla CN, McDonnell WJ, Ram R, et al. (2021) Single-cell analysis shows that adipose tissue of persons with both HIV and diabetes is enriched for clonal, cytotoxic, and CMV-specific CD4+ T cells. Cell Rep Med 2:100205. 10.1016/j.xcrm.2021.100205.

[28] Picelli S, Faridani OR, Björklund ÅK, Winberg G, Sagasser S and Sandberg R (2014) Full-length RNA-seq from single cells using Smart-seq2. Nature Protocols 9:171–181. 10.1038/nprot.2014.006.

[29] Jaitin DA, Kenigsberg E, Keren-Shaul H, et al. (2014) Massively parallel single-cell RNA-seq for marker-free decomposition of tissues into cell types. Science (New York, N.Y.) 343:776–779. 10.1126/science.1247651.

[30] Islam S, Zeisel A, Joost S, La Manno G, Zajac P, Kasper M, Lönnerberg P and Linnarsson S (2014) Quantitative single-cell RNA-seq with unique molecular identifiers. Nature Methods 11:163–166. 10.1038/nmeth.2772.

[31] Kivioja T, Vähärautio A, Karlsson K, Bonke M, Enge M, Linnarsson S and Taipale J (2012) Counting absolute numbers of molecules using unique molecular identifiers. Nature Methods 9:72–74. 10.1038/nmeth.1778.

[32] Grün D, Kester L and van Oudenaarden A (2014) Validation of noise models for single-cell transcriptomics. Nature Methods 11:637–640. 10.1038/nmeth.2930.

[33] Butler A, Hoffman P, Smibert P, Papalexi E and Satija R (2018) Integrating single-cell transcriptomic data across different conditions, technologies, and species. Nature Biotechnology 36:411–420. 10.1038/nbt.4096.

[34] Zhang Y, Parmigiani G and Johnson WE (2020) ComBat-seq: batch effect adjustment for RNA-seq count data. NAR Genomics and Bioinformatics 2:lqaa078. 10.1093/nargab/lqaa078.

[35] Chen EY, Tan CM, Kou Y, Duan Q, Wang Z, Meirelles GV, Clark NR and Ma’ayan A (2013) Enrichr: interactive and collaborative HTML5 gene list enrichment analysis tool. BMC Bioinformatics 14:128. 10.1186/1471-2105-14-128.

[36] Supek F, Bošnjak M, škunca N and šmuc T (2011) REVIGO Summarizes and Visualizes Long Lists of Gene Ontology Terms. PLOS ONE 6:e21800. 10.1371/journal.pone.0021800.

[37] Zappia L and Oshlack A (2018) Clustering trees: a visualization for evaluating clusterings at multiple resolutions. Gigascience 7:10.1093/gigascience/giy083.

[38] Pfeiffer-Guglielmi B, Fleckenstein B, Jung G and Hamprecht B (2003) Immunocytochemical localization of glycogen phosphorylase isozymes in rat nervous tissues by using isozyme-specific antibodies. J Neurochem 85:73–81. 10.1046/j.1471-4159.2003.01644.x.

[39] Becht E, McInnes L, Healy J, Dutertre C-A, Kwok IWH, Ng LG, Ginhoux F and Newell EW (2019) Dimensionality reduction for visualizing single-cell data using UMAP. Nature Biotechnology 37:38–44. 10.1038/nbt.4314.

[40] del Amo EM, Vellonen K-S, Kidron H and Urtti A (2015) Intravitreal clearance and volume of distribution of compounds in rabbits: In silico prediction and pharmacokinetic simulations for drug development. European Journal of Pharmaceutics and Biopharmaceutics 95:215–226. https://doi.org/10.1016/j.ejpb.2015.01.003.

[41] Raeisossadati R, Ferrari MFR, Kihara AH, AlDiri I and Gross JM (2021) Epigenetic regulation of retinal development. Epigenetics Chromatin 14:11. 10.1186/s13072-021-00384-w.

[42] Mo A, Luo C, Davis FP, et al. (2016) Epigenomic landscapes of retinal rods and cones. eLife 5:e11613. 10.7554/eLife.11613.

[43] Hughes AEO, Enright JM, Myers CA, Shen SQ and Corbo JC (2017) Cell Type-Specific Epigenomic Analysis Reveals a Uniquely Closed Chromatin Architecture in Mouse Rod Photoreceptors. Scientific Reports 7:43184. 10.1038/srep43184.

[44] Ueno K, Iwagawa T, Kuribayashi H, Baba Y, Nakauchi H, Murakami A, Nagasaki M, Suzuki Y and Watanabe S (2016) Transition of differential histone H3 methylation in photoreceptors and other retinal cells during retinal differentiation. Scientific Reports 6:29264. 10.1038/srep29264.

[45] Burgoyne LA (1999) The mechanisms of pyknosis: hypercondensation and death. Exp Cell Res 248:214–22. 10.1006/excr.1999.4406.

[46] Basavarajappa BS and Subbanna S (2021) Histone Methylation Regulation in Neurodegenerative Disorders. International journal of molecular sciences 22:4654. 10.3390/ijms22094654.

[47] Zhao M, Tao Y and Peng G-H (2020) The Role of Histone Acetyltransferases and Histone Deacetylases in Photoreceptor Differentiation and Degeneration. Int J Med Sci 17:1307–1314. 10.7150/ijms.43140.

[48] Marinova Z, Leng Y, Leeds P and Chuang D-M (2011) Histone deacetylase inhibition alters histone methylation associated with heat shock protein 70 promoter modifications in astrocytes and neurons. Neuropharmacology 60:1109–1115. 10.1016/j.neuropharm.2010.09.022.

[49] Halsall JA, Turan N, Wiersma M and Turner BM (2015) Cells adapt to the epigenomic disruption caused by histone deacetylase inhibitors through a coordinated, chromatin-mediated transcriptional response. Epigenetics Chromatin 8:29–29. 10.1186/s13072-015-0021-9.

[50] Heinemann B, Nielsen JM, Hudlebusch HR, et al. (2014) Inhibition of demethylases by GSK-J1/J4. Nature 514:E1–E2. 10.1038/nature13688.

[51] Mu M-D, Qian Z-M, Yang S-X, Rong K-L, Yung W-H and Ke Y (2020) Therapeutic effect of a histone demethylase inhibitor in Parkinson’s disease. Cell Death & Disease 11:927. 10.1038/s41419-020-03105-5.

[52] Cenik BK and Shilatifard A (2021) COMPASS and SWI/SNF complexes in development and disease. Nature Reviews Genetics 22:38–58. 10.1038/s41576-020-0278-0.

[53] Bochyńska A, Lüscher-Firzlaff J and Lüscher B (2018) Modes of Interaction of KMT2 Histone H3 Lysine 4 Methyltransferase/COMPASS Complexes with Chromatin. Cells 7:17. 10.3390/cells7030017.

[54] Qi L, Tsai B and Arvan P (2017) New Insights into the Physiological Role of Endoplasmic Reticulum-Associated Degradation. Trends Cell Biol 27:430–440. 10.1016/j.tcb.2016.12.002.

[55] Kroeger H, Chiang W-C and Lin JH (2012) Endoplasmic reticulum-associated degradation (ERAD) of misfolded glycoproteins and mutant P23H rhodopsin in photoreceptor cells. Advances in experimental medicine and biology 723:559–565. 10.1007/978-1-4614-0631-0_71.

[56] Chatterjee N and Walker GC (2017) Mechanisms of DNA damage, repair, and mutagenesis. Environ Mol Mutagen 58:235–263. 10.1002/em.22087.

[57] Okita A, Murakami Y, Shimokawa S, et al. (2020) Changes of Serum Inflammatory Molecules and Their Relationships with Visual Function in Retinitis Pigmentosa. Investigative Ophthalmology & Visual Science 61:30–30. 10.1167/iovs.61.11.30.

[58] Lefevere E, Toft-Kehler AK, Vohra R, Kolko M, Moons L and Van Hove I (2017) Mitochondrial dysfunction underlying outer retinal diseases. Mitochondrion 36:66–76. https://doi.org/10.1016/j.mito.2017.03.006.

[59] Jiang X and Wang X (2004) Cytochrome C-Mediated Apoptosis. Annual Review of Biochemistry 73:87–106. 10.1146/annurev.biochem.73.011303.073706.

[60] Huang H, Li F, Alvarez RA, Ash JD and Anderson RE (2004) Downregulation of ATP Synthase Subunit-6, Cytochrome c Oxidase-III, and NADH Dehydrogenase-3 by Bright Cyclic Light in the Rat Retina. Investigative Ophthalmology & Visual Science 45:2489–2496. 10.1167/iovs.03-1081.

[61] Usui S, Komeima K, Lee SY, et al. (2009) Increased Expression of Catalase and Superoxide Dismutase 2 Reduces Cone Cell Death in Retinitis Pigmentosa. Molecular Therapy 17:778–786. https://doi.org/10.1038/mt.2009.47.

[62] Usui S, Oveson BC, Iwase T, et al. (2011) Overexpression of SOD in retina: Need for increase in H2O2-detoxifying enzyme in same cellular compartment. Free Radical Biology and Medicine 51:1347–1354. https://doi.org/10.1016/j.freeradbiomed.2011.06.010.

[63] Rozpedek W, Pytel D, Mucha B, Leszczynska H, Diehl JA and Majsterek I (2016) The Role of the PERK/eIF2α/ATF4/CHOP Signaling Pathway in Tumor Progression During Endoplasmic Reticulum Stress. Curr Mol Med 16:533–544. 10.2174/1566524016666160523143937.

[64] Bhootada Y, Kotla P, Zolotukhin S, Gorbatyuk O, Bebok Z, Athar M and Gorbatyuk M (2016) Limited ATF4 Expression in Degenerating Retinas with Ongoing ER Stress Promotes Photoreceptor Survival in a Mouse Model of Autosomal Dominant Retinitis Pigmentosa. PLOS ONE 11:e0154779. 10.1371/journal.pone.0154779.

[65] Chu X, Zhong L, Yu L, et al. (2020) GSK-J4 induces cell cycle arrest and apoptosis via ER stress and the synergism between GSK-J4 and decitabine in acute myeloid leukemia KG-1a cells. Cancer Cell International 20:209. 10.1186/s12935-020-01297-6.

[66] Cideciyan AV, Jacobson SG, Beltran WA, et al. (2013) Human retinal gene therapy for Leber congenital amaurosis shows advancing retinal degeneration despite enduring visual improvement. Proc Natl Acad Sci U S A 110:E517–25. 10.1073/pnas.1218933110.

