## Supplementary Figures and Supplementary Table S1 for "Increased H3K27 trimethylation contributes to cone survival in a mouse model of cone dystrophy"

**Table S1. Summary of primary and secondary antibodies used for immunohistochemistry**

| **Antibody** | **Host species** | **Supplier** | **Cat no.** | **Working dilution** | **Incubation** |
|---|---|---|---|---|---|
| GFP Polyclonal Antibody, AlexaFluor 488 | Rabbit | Invitrogen | A-21311 | 1:500 | 2 hours room temperature |
| Glial Fibrillary Acidic Protein Polyclonal Antibody | Rabbit | DAKO | Z0334 | 1:500 | 4ºC overnight |
| Tri-Methyl-Histone H3 (Lys27) | Rabbit | Cell Signalling | 9733S | 1:200 | 4ºC overnight |
| Histone H3 [Trimethyl Lys9] Antibody | Mouse | Novus Biologicals | NBP1-30141SS | 1:200 | 4ºC overnight |
| Anti-Opsin Antibody, Red/Green | Rabbit | Millipore | AB5405 | 1:1000 | 4ºC overnight |
| Anti-Opsin Antibody, Blue | Rabbit | Millipore | AB5407 | 1:1000 | 4ºC overnight |
| Glycogen Phosphorylase | Guineapig | Custom made [1] | N/A | 1:1000 | 4ºC overnight |
| Anti-Cone Arrestin Antibody | Rabbit | Millipore | AB15282 | 1:1000 | 4ºC overnight |
| Goat Anti-Rabbit IgG H+L AlexaFluor 568 | Rabbit | Abcam | AB175471 | 1:500 | 2 hours room temperature |
| Goat Anti-Mouse IgG H+L AlexaFluor 568 | Mouse | ThermoFisher | AB11031 | 1:500 | 1 hour room temperature |
| Goat Anti-Guineapig IgG FITC | Guineapig | Merck | F6261 | 1:300 | 1 hour room temperature |

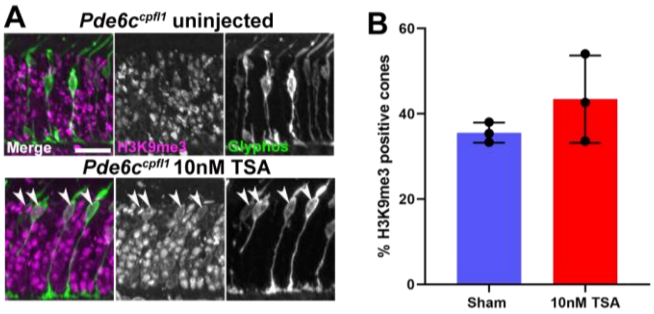


**Fig. S1 A** H3K9me3 staining appeared to be increased in the cones after the HDAC inhibitor TSA was administered intravitreally in *Pde6c$^{cpfl1}$* mice. H3K9me3 (magenta) and cone (stained in green with Glyphos antibody) co-localization denoted by arrows. **B** Quantification of the percentage of H3K9me3 positive cones in sham controls and TSA-treated *Pde6c$^{cpfl1}$* mice. Welsh's T-test, n=3, *P*>0.05

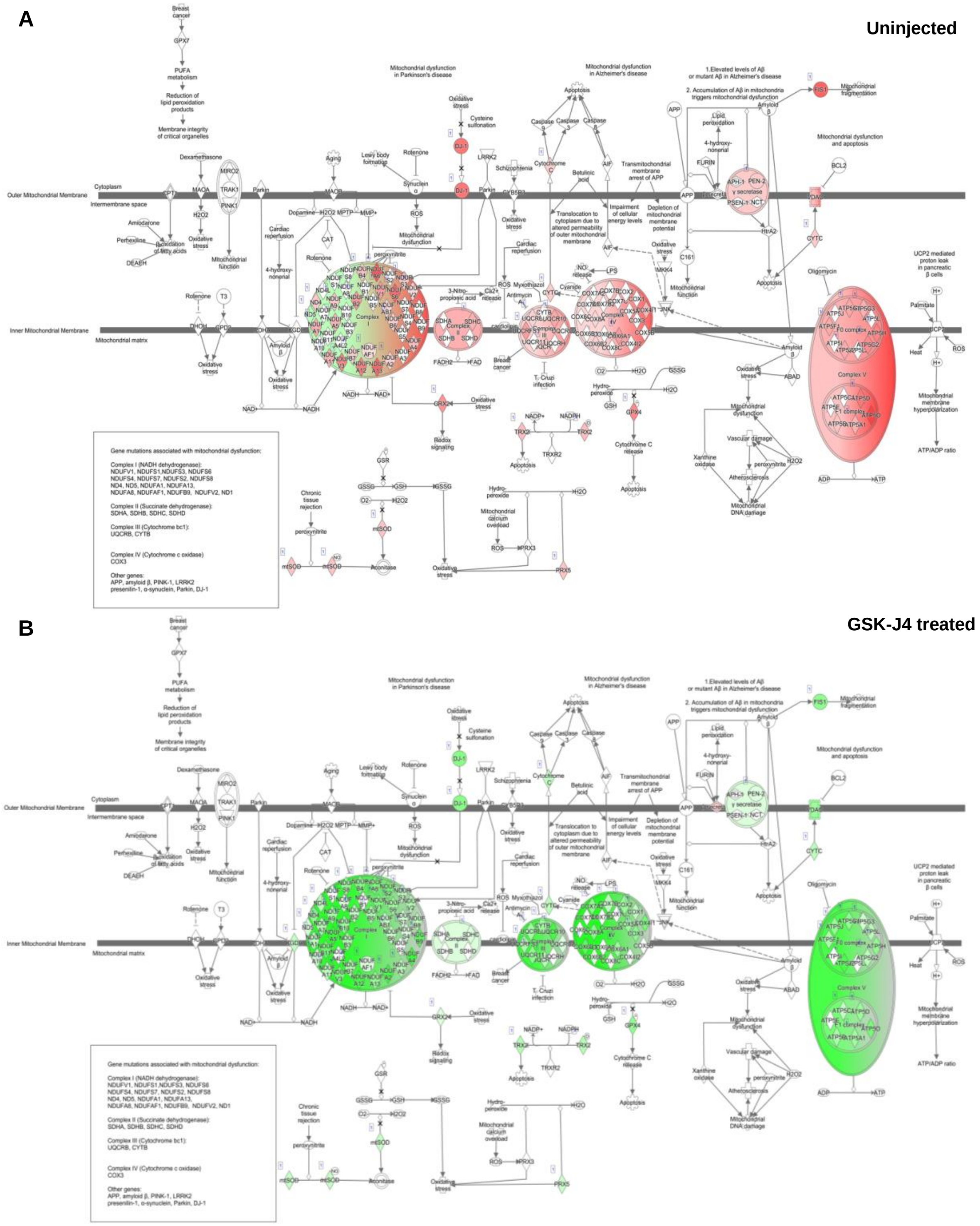


**Fig. S2** Schematic diagrams of the mitochondrial dysfunction pathway in **A** uninjected and **B** GSK-J4 treated *Pde6c*.GFP cones, with expression levels of genes overlaid. After treatment with GSK-J4, we observed a significant change in the expression profile, with downregulation of key genes in complexes I-V, which are responsible for oxidative phosphorylation. Diagram downloaded from QIAGEN Ingenuity Pathway Analysis, genes shown in the diagram have >0.3 absolute log fold change, and an unadjusted $P$<0.01. Green downregulation; red upregulation

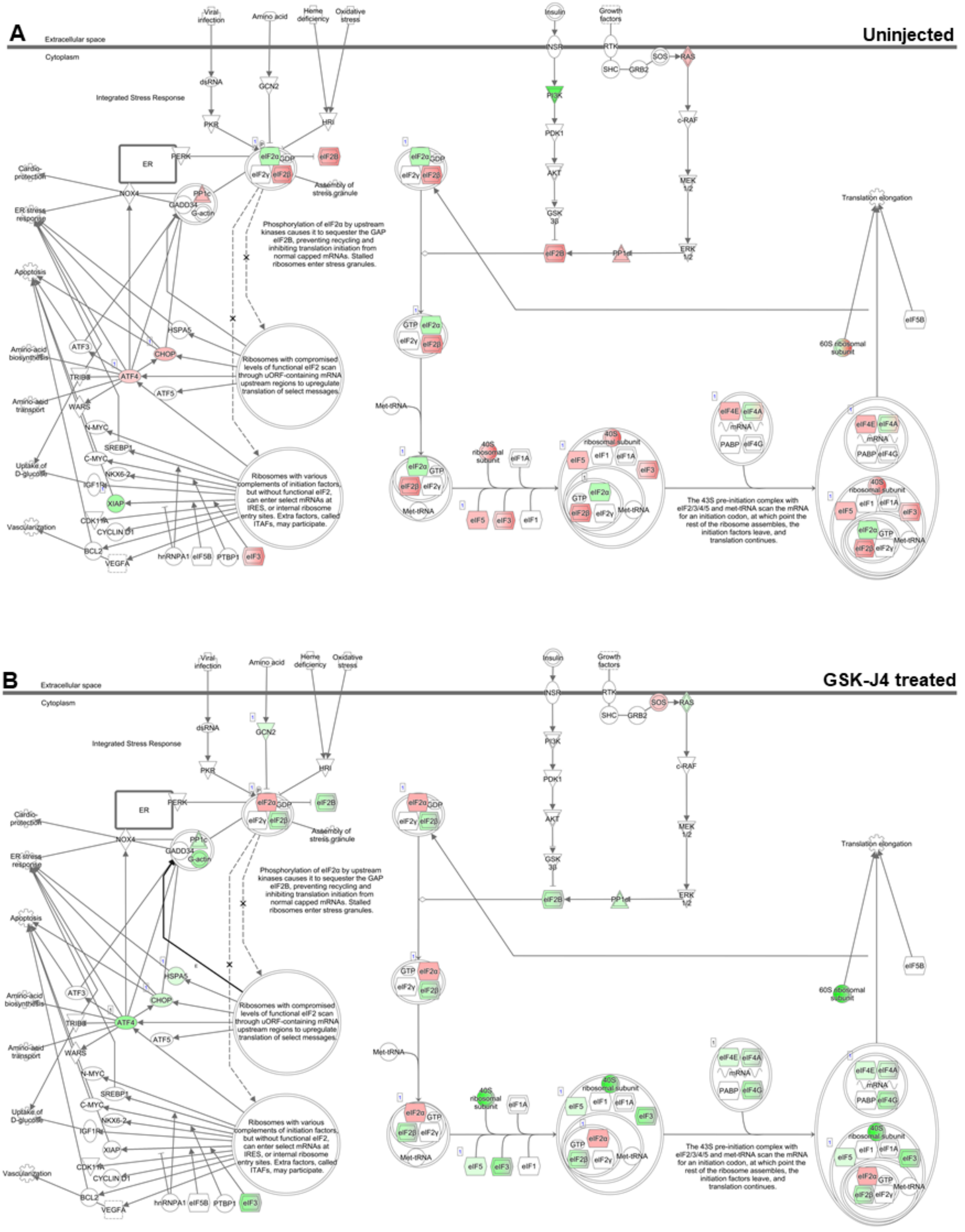


**Fig. S3** Schematic diagrams of the EIF2 signaling pathway in **A** uninjected and **B** GSK-J4 treated *Pde6c*.GFP cones, with expression levels of genes overlaid. After treatment with GSK-J4, we noted downregulation of the CHOP cell death pathway, and an overall reduction in protein production. Diagram downloaded from QIAGEN Ingenuity Pathway Analysis, genes shown in the diagram have >0.3 absolute log fold change, and an unadjusted *P*<0.01. Green downregulation; red upregulation

[1] Pfeiffer-Guglielmi B, Fleckenstein B, Jung G and Hamprecht B (2003) Immunocytochemical localization of glycogen phosphorylase isozymes in rat nervous tissues by using isozyme-specific antibodies. J Neurochem 85:73-81. 10.1046/j.1471-4159.2003.01644.x.
